# Optimized SMRT-UMI protocol produces highly accurate sequence datasets from diverse populations – application to HIV-1 quasispecies

**DOI:** 10.1101/2023.02.23.529831

**Authors:** Dylan H. Westfall, Wenjie Deng, Alec Pankow, Hugh Murrell, Lennie Chen, Hong Zhao, Carolyn Williamson, Morgane Rolland, Ben Murrell, James I. Mullins

## Abstract

Pathogen diversity resulting in quasispecies can enable persistence and adaptation to host defenses and therapies. However, accurate quasispecies characterization can be impeded by errors introduced during sample handling and sequencing which can require extensive optimizations to overcome. We present complete laboratory and bioinformatics workflows to overcome many of these hurdles. The Pacific Biosciences <u>s</u>ingle <u>m</u>olecule <u>r</u>eal-<u>t</u>ime platform was used to sequence PCR amplicons derived from cDNA templates tagged with <u>u</u>niversal <u>m</u>olecular identifiers (SMRT-UMI). Optimized laboratory protocols were developed through extensive testing of different sample preparation conditions to minimize between-template recombination during PCR and the use of UMI allowed accurate template quantitation as well as removal of point mutations introduced during PCR and sequencing to produce a highly accurate consensus sequence from each template. Handling of the large datasets produced from SMRT-UMI sequencing was facilitated by a novel bioinformatic pipeline, Probabilistic Offspring Resolver for Primer IDs (PORPIDpipeline), that automatically filters and parses reads by sample, identifies and discards reads with UMIs likely created from PCR and sequencing errors, generates consensus sequences, checks for contamination within the dataset, and removes any sequence with evidence of PCR recombination or early cycle PCR errors, resulting in highly accurate sequence datasets. The optimized SMRT-UMI sequencing method presented here represents a highly adaptable and established starting point for accurate sequencing of diverse pathogens. These methods are illustrated through characterization of human immunodeficiency virus (HIV) quasispecies.

**Author Summary:** There is a great need to understand the genetic diversity of pathogens in an accurate and timely manner, but many errors can be introduced during the sample handling and sequencing steps which may prevent accurate analyses. In some cases, the errors introduced during these steps can be indistinguishable from real genetic variation and prevent analyses from identifying true sequence variation present in the pathogen population. There are established methods which can help to prevent these types of errors, but can involve many different steps and variables, all of which must be optimized and tested together to ensure the desired effect. Here we show results from testing different methods on a set of HIV+ blood plasma samples and arrive at a streamlined laboratory protocol and bioinformatic pipeline which prevents or corrects for different types of errors that can arise in sequence datasets. These methods should be an accessible starting point for anyone wanting accurate sequencing without extensive optimizations.

## Introduction

Viral quasispecies are populations of genetically related non-identical viruses – typically RNA viruses – with the potential to facilitate adaptation to a changing host environment and thus contribute features beneficial to the survival of the overall population within a host[1-3]. Viral quasispecies have traditionally been examined by Sanger sequencing of individual viral templates, yet such an approach is low throughput (usually limited to a few dozen viruses) and cost-intensive, as limiting dilution is required for highly accurate sequence derivation[4]. While next-generation sequencing (NGS) provides a high throughput alternative, the short read lengths, e.g., using the Illumina platform, typically prevent studies of linkage across genes much longer than sequence reads (150-600 bp)[5, 6]. Pacific Biosciences (PacBio) single-molecule real-time (SMRT)[7] and nanopore[8] sequencing technologies are both 3^rd^ generation long-read, high-throughput sequencing platforms capable of producing read lengths in excess of 10 Kb[9]. However, errors inherent in these technologies and introduced during PCR can persist within the population of sequenced molecules and prevent accurate assessment of viral quasispecies. Furthermore, PCR-mediated recombination (chimera formation)[10] can erroneously link viral mutations and be mistaken for viral recombination *in vivo*, thus obscuring accurate quasispecies assessment.

Incorporation of molecular tags composed of random sequences (typically 6-12 nt), referred to as primer IDs[11] or unique molecular identifiers (UMIs)[12], into cDNA primers used in the first step of RNA virus genome amplification tags every cDNA molecule with a random sequence that is preserved during subsequent PCR amplification. Importantly, the number of cDNA templates that are then amplified can be accurately determined by the number of unique UMIs recovered[7]. By collecting sequencing reads with the same UMI (hereafter referred to as UMI families), then generating a consensus, PCR and sequencing errors can be removed to accurately estimate the sequence of each original cDNA template. A single UMI (sUMI) alone cannot always identify products that incurred template switching during PCR (often referred to as recombination). However, incorporating a second UMI on the opposite strand labels each viral template with a unique combination of UMI sequences, one on either end of the molecule (duplex UMI or dUMI)[13]. By comparing the frequencies of different UMI combinations and comparing UMI sequences between families, reads representing molecules that “recombined” during PCR can be identified. However, drawbacks to the dUMI approach, such as loss of sample material during multiple purification steps necessary to remove excess primers, and fixation of errors during second strand synthesis led us to seek sample preparation conditions resulting in low levels of recombination such that sUMI methods give accuracy approximately equivalent to dUMI. dUMI labeled samples were thus prepared varying different sample preparation methods thought to influence error rates such as template limit per PCR, a size selection step after 1^st^ round of PCR, and PCR cycle number.

The PacBio SMRT platform was chosen for sequencing because of the lower error rate derived from circular consensus sequences (CCS)[14] compared to current nanopore approaches[15, 16]. As part of our workflow, bioinformatics pipelines were developed to take CCS as input and produce tables of all observed UMI combinations as well as consensus sequences prepared from the sUMI or dUMI read subsets of each family. This allowed for direct comparisons between consensus sequences from sUMI and dUMI methods. The core computational operations are implemented in the Julia language for scientific computing and packaged into the Snakemake workflow management system to perform a series of tasks such as demultiplexing, UMI calling, consensus generation, and quality filtering based on agreement between reads.

Comparing UMI and sequence results from different preparation methods identified conditions which limited PCR recombination and where sequences from sUMI and dUMI methods were identical. This allowed use of only a single UMI while maintaining the accuracy of a dual UMI preparation.

After optimizing laboratory methods for only sUMI, an improved pipeline, named PORPIDpipeline, was created with additional functionality. In brief, the pipeline filters by read length and quality, separates sequences by sample ID and then by UMI, and removes: UMI families likely to be “offspring” generated by errors from real UMI families; UMI families characterized as heteroduplexes; UMIs not matching the expected length; UMI families with fewer than 5 reads. Consensus sequences are generated for each remaining UMI family, and sequences with consensus agreement less than 0.7 at any base position are discarded. Downstream processing includes alignment and trimming against reference panels, identifying likely contamination, both from common lab strains and cross-contamination between samples prepared together, and providing HTML reports with various quantity and quality metrics, as well as visualizations of phylogenies, alignments, APOBEC-mediated hypermutation, and more.

The optimized sUMI workflow combined with the PORPIDpipeline represents a streamlined and generalized sequencing approach which can be highly adaptable and is anticipated to help advance the study of pathogens that develop intra-host genetic diversity during infection as well as studies of diverse organism communities in general.

## Results

### Optimization of Laboratory Protocols

Fig. 1 presents the overall workflow using either sUMI and dUMI methods, both of which consists of primer design, template preparation, SMRT sequencing and data processing, identification of CCS reads with the same UMI, and derivation of consensus sequences for each viral template. As shown in Fig. 1A, UMIs can either be applied to cDNA alone (single or sUMI) or to both first strand cDNA and second strand synthesis products (dual or dUMI). Fig. 1B includes the initial reverse transcription step of cDNA synthesis from RNA templates; errors introduced at this step cannot be corrected at any point downstream. cDNA synthesis errors accumulate according to the fidelity of the chosen reverse transcriptase and the relevant reaction conditions (e.g., reported error rates for MMLV-based reverse transcriptases range from one per 15.5 kb synthesized[17] to one per 27 kb synthesized[18]). cDNA products are purified following the cDNA synthesis step to remove unincorporated primers (Fig. 1B). In the case of dUMI, the purified cDNA is subjected to second strand synthesis with a primer containing a second UMI (steps shaded in gray in Fig. 1B). The resulting dsDNA product is again purified to remove unincorporated primers. When only sUMI are to be employed, or after second strand synthesis in the dUMI protocol, the purified products are subjected to nested PCR, in order to generate sufficient material for addition of SMRT bells[19] and sequencing. As heteroduplexes formed in the last cycle of PCR can result in reads containing artifactual UMI sequences, their formation is avoided by performing a single cycle of synthesis and extension (Final Extension) using fresh reagents to bias utilization of PCR primers as opposed to incomplete PCR products serving as primers[20]. Without this step, many reads will fail the heteroduplex filter in the first steps of the pipeline and be discarded, leading to lower read depths for each UMI family.

**Figure 1.**
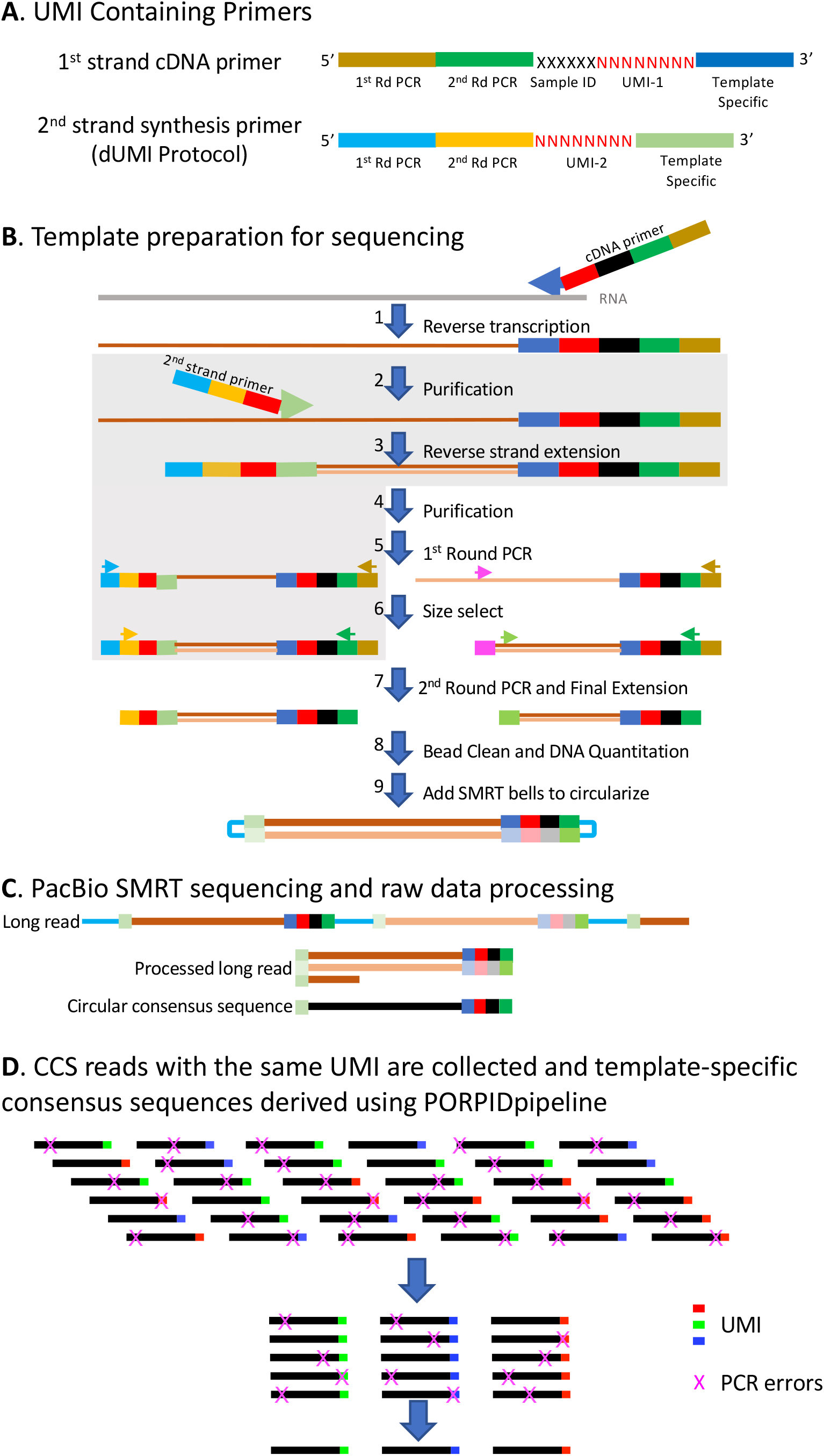
Overview of the viral genome sequencing process. **A**. cDNA synthesis is conducted using a library of primers with the following structure (reading from the 5’ end of the primer): Universal reverse 1st and 2nd PCR primers, followed by a 6 nt sample ID, followed by an 8 nt random sequence corresponding to the universal molecular identifier (UMI) and then a viral template-specific sequence at the 3’ end. The 2nd strand synthesis primer is identical except that it lacks the 6 nt sample ID. **B**. Overview of the sequencing template generation process. Following cDNA synthesis (1), unused primers are removed by bead cleaning (2/4). For sUMI samples this step (4) is followed by 1st round PCR (5). For dUMI samples (indicated by the grey box), a second strand primer containing another UMI is introduced by a single round of PCR (3), followed by another round of bead cleaning to remove unused primers (4). The 1st round of PCR is conducted using the 1st round universal reverse primer described in panel a, and either a forward HIV template-specific primer (magenta) for sUMI or a universal forward primer for dUMI (light blue) (5). PCR products below the expected size are then removed using bead purification or the Blue Pippin instrument (6) and a 2nd round of PCR followed by a single cycle final extension with new reagents and conducted with inner universal reverse and HIV-specific forward primers for sUMI (green) or inner universal forward (yellow) for dUMI (7). PCR primers are removed by bead cleaning and the products quantified (Qubit) and reactions pooled (8). SMRTbell barcoded adapters are ligated onto the ends of the PCR fragments to create a circular molecule (9). **C**. Illustration of PacBio Sequence Generation. A sequencing primer is annealed to the SMRTbell template and DNA polymerase is bound to the complex. This polymerase-amplicon-adaptor complex is then loaded into zero-mode waveguides (ZMWs) where replication occurs, producing nucleotide-specific fluorescence upon incorporation into DNA. The polymerase repeatedly replicates the circularized strand, producing one long read with randomly distributed errors. Post-run, the sequences are processed -SMRTbell sequences are bioinformatically trimmed away, single-molecule fragments are aligned, and a circular consensus sequence (CCS) is generated. **D**. The final step in sequence generation for sUMI samples is the identification of CCS with the same UMI and generation of a template-specific consensus sequence using the PORPIDpipeline.

By applying circular consensus sequencing[19] (Fig. 1C) to improve the accuracy of SMRT sequencing and retaining only reads with a Quality (Q) score ≥ 20, long-range (>10 Kb) consensus reads with an average accuracy of 99.8% have recently been reported[14]. In our optimized workflow, a consensus of sUMI labelled templates is generated (Fig. 1D) to remove additional PCR errors. While prior PacBio instruments exhibited loading bias, increasing the frequency of shorter amplicons, with current Sequel II instrument reagents and software, roughly equivalent CCS recovery of 2.5 kb to 6 kb amplicons were obtained when loaded onto the same plate at equimolar ratios.

### dUMI sequencing

dUMI can also be used and a consensus sequence derived from each family of reads with the same combination of UMI sequences. Use of dUMI and the requirement for consistent UMI on each strand to perform consensus generation, lowers the error rate to approximately 10^−9^/base incorporated in steps beyond creation of the cDNA[13]. A second advantage of using dUMI is that recombinant molecules are readily identified by detection of discordant dUMI and excluded; recombinant molecules are produced by template switching when incomplete products act as primers during the PCR. Table 1 illustrates a scenario in which each member of a family of CCS reads shares the same UMI-1 sequence (applied during first strand cDNA synthesis) yet the UMI-2 sequences (applied to the opposite end of the molecule during second strand synthesis of the dUMI protocol) differ across the reads. In this case, the most prevalent dUMI combination (Rank 1) is inferred to represent the original template, and less frequent combinations are inferred to represent combinations introduced by recombination, or PCR or sequencing errors accumulating within the UMI-2 sequence.

**Table 1.** Family of reads sharing same UMI-1 sequence (sUMI family)

| UMI-1 | UMI-2 | Family Size | Rank | Levenshtein Distance* | Matches UMI-2 from different family |
| --- | --- | --- | --- | --- | --- |
| CCGTGGAT | CAATCACC | 180 | 1 |  |  |
| CCGTGGAT | CAATCCACC | 1 | 2 | 1 | No |
| CCGTGGAT | AATTCACC | 1 | 3 | 2 | No |
| CCGTGGAT | AATGATGT | 1 | 4 | 5 | No |
| CCGTGGAT | GGGAACCC | 1 | 5 | 5 | Yes |
| CCGTGGAT | GTGGTACT | 1 | 6 | 6 | No |
| CCGTGGAT | TAAGATGG | 1 | 7 | 6 | Yes |

\* Distance from rank 1 UMI-2 to other UMI-2 sequences observed.

### Optimization of the sUMI protocol

One disadvantage of the dUMI approach, specific to the use of single stranded templates such as cDNA, is that errors generated during 2^nd^ strand synthesis would be fixed, as only the molecules with the second strand synthesis primer would be amplified in subsequent cycles. A second cDNA-specific disadvantage is that significant material loss can occur during bead purification to remove primers, which is required after both first and second strand synthesis reactions. Sample preparation conditions were therefore sought to minimize the rate of recombination to allow use of the sUMI workflow without specific identification of recombinant reads via detection of discordant dUMI. To assess different reaction conditions, samples were prepared for sequencing using the dUMI approach with the goal of sequencing 100 total cDNA templates. The number of estimated input templates (e.g., 25, 50, 100) per PCR, purification of the 1^st^ round PCR, and PCR cycle number were varied to test different combinations. First round PCR reactions from each sample were divided and subjected to size selection with either AMPure XP beads (Beckman Coulter, Indianapolis, IN) or the Blue Pippin gel electrophoresis instrument (Sage Science, Inc., Beverly, MA) and compared to fractions that were subjected to identical dilutions but not purified. These separate preparations were amplified in a twenty-two cycle 2^nd^ round PCR. To test whether lower PCR cycle numbers at high template input could help to reduce recombination rates, the Blue Pippin purified fractions from the 100 template reactions were separately amplified in 2^nd^ round PCRs consisting of 20 cycles. All 2^nd^ round preparations were then carried forward in an identical manner for subsequent purification, PCR, addition of the Index primers used for demultiplexing, SMRT sequencing, and analysis.

For the studies described here, ~3kb amplicons were derived from virion-associated HIV RNA present in blood plasma from a total of 33 different sample preparations (S1 Table). These studies yielded 6,666 sUMI families that passed all UMI filtering steps of the pipeline and hence were deemed “likely real”. Next, examining the UMI-2 tags found in these sUMI families resulted in identification of 63,139 unique dUMI families. The origin of the large increase in dUMI compared to sUMI families is illustrated in Table 1, where reads with the same UMI-1 (sUMI family) are found to contain seven different UMI-2 sequences (with each combination thus representing a distinct dUMI family). Levenshtein distances[21] (LDs) were calculated for every distinct UMI-2 (contained within the 2^nd^ strand synthesis primer) sharing the same UMI-1 (contained within the cDNA primer). These distances count the fewest number of substitutions, insertions, or deletions necessary to create each observed UMI-2, relative to the most prevalent (rank 1) UMI-2 in that family (e.g., Table 1). dUMI combinations with distances of 1 or possibly 2 are more likely to result from PCR or sequencing errors, as the risk of accumulating greater than 2 errors via PCR or sequencing within the relatively short 8 bp UMI region is very low and UMI-2 heteroduplexes should be rare after filtering.

To identify which UMI combinations within each sUMI family were the result of recombination during PCR, the rank >1 UMI-2 sequences from each family were compared to the rank 1 UMI-2 sequences from all other families in the same sample. Conservatively, any rank >1 families with a UMI-2 that matched a rank 1 family were judged to result from recombination. Nearly all rank 1 families were found to donate both UMI-1 and UMI-2 sequences to recombinant reads. In most cases, the size of the recombinant family was 1 while the size of the rank 1 families the recombinant was derived from were much larger (Fig. 2A); the median size of rank 1 families that donated a UMI sequence to a recombinant was 43, while the median of those that did not was 9.

**Figure 2.**
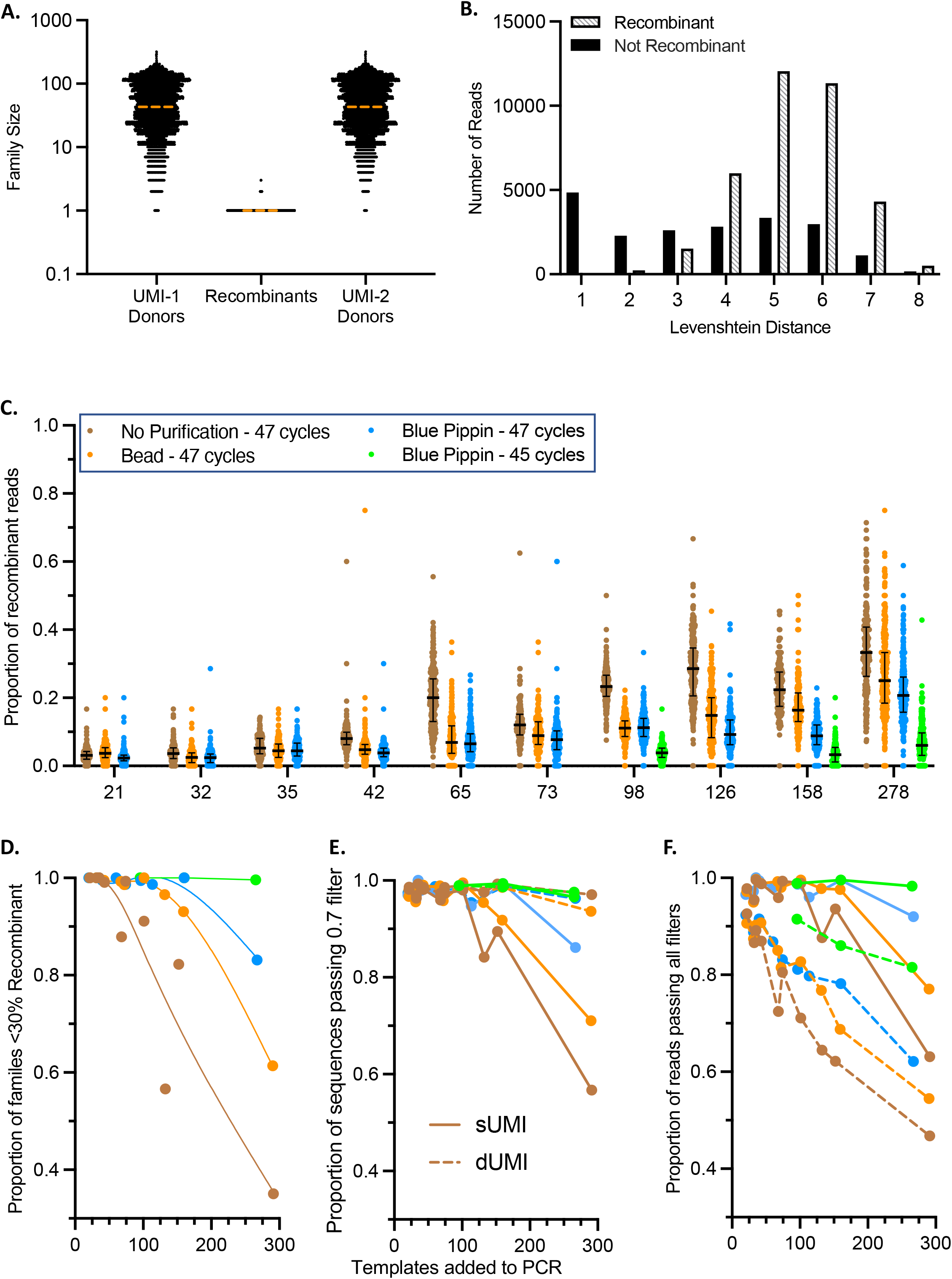
Identification of recombinant reads and conditions that limit their effect. **A**. Family size is indicated for each recombinant family as well as all rank 1 families the recombinants received UMI-1 or UMI-2 sequences from. The dashed horizontal orange lines indicate the medians. **B**. Levenshtein distances (LD) are assigned to each read in a family. For each LD, counts of reads judged to be recombinants or not are indicated. **C**. Proportion of recombinant reads for each sUMI family, grouped by average template input into PCR, where each dot represents an individual sUMI family, colored by purification method and total PCR cycle number. **D**. The proportion of sUMI families in each sample containing fewer than 30% recombinant reads at different template inputs into PCR. **E**. Proportion of sequences that pass the 0.7 minimum agreement filter in the PORPIDpipeline at different template inputs into PCR. **F**. Proportion of reads passing all filters in the PORPIDpipeline at different template inputs into PCR.

As anticipated, the UMI-2 sequences most similar to the rank 1 family (LD 1 or 2) did not frequently match any other family. In contrast, ~80% of the UMI families with LD >3 did match other families (Fig. 2B). We judged that it was unlikely that many of these resulted from an accumulation of 4+ changes to the original UMI-2 sequence, but more likely these are the result of a recombination event. As we required an exact match between UMI-2 sequences to classify a family as recombinant, the numbers of families noted as recombinants are likely an underestimation, as any PCR error in the UMI-2 after a recombination event would appear as a unique sequence.

Next, PCR input template number and purification of 1^st^ round PCR products were optimized to limit recombinant yield. While we aimed to add 25, 50, or 100 templates to each PCR based on estimates from limiting dilution PCR, the number varied from 21-278 (S1 Table). Two purification methods were compared to unpurified PCR products – size selection using the Blue Pippin instrument and AMPure XP beads. Smaller, off-target PCR products are frequently generated when amplifying large amplicons. Prevalence of these off-target products varies by sample and PCR conditions but without effective removal can prevent efficient amplification and recovery of larger, targeted amplicons. Additionally, off-target products present in the final sequencing mixture can reduce sequencing depth of the target amplicon. By purifying the 1^st^ round PCR products, some off-target products are removed, and subsequent PCR rounds efficiently amplify the remaining PCR products. When evaluating and selecting a purification method, a somewhat wide range of molecular sizes should nonetheless be captured to avoid loss of molecules of interest that may have suffered large internal deletions or insertions in vivo.

Blue Pippin purification resulted in substantially more efficient removal of smaller PCR products and somewhat greater yield of the desired product than purification with beads compared to unpurified fractions (Fig. S1A). Fig. 2C shows the proportion of recombinant reads found in each sUMI family as a function of the number of templates in each PCR and the PCR product purification method. At lower input numbers (≤ 50), relatively low fractions of recombinant reads were observed without purification. Increasing initial template number per PCR resulted in benefits from purification, with Blue Pippin having generally greater impact compared to bead purification. Furthermore, reduction of PCR cycles reduced recombination in the Blue Pippin fraction to greater effect, resulting in rates comparable to much lower template inputs. This indicates that using the fewest PCR cycles that still result in sufficient DNA yield for sequencing is ideal. This can be achieved by running multiple parallel PCR reactions. While unpurified samples generally have a greater proportion of recombinant reads, target bands on 1% agarose gels are stronger in purified samples using the same cycle number (Fig. S1A), suggesting recombination rates are influenced substantially by the prevalence of smaller PCR products in the mixture. Greater sequence numbers can be also obtained by increasing the number of PCR reactions at template inputs that prevent high recombination rates.

To facilitate the use of the sUMI workflow, strict methods were developed within the bioinformatics pipeline to remove families with sequence variation great enough to impact the consensus. A sUMI family is discarded if the minimum agreement at any position in its alignment was <70% (hereafter referred to as the 0.7 filter). This removes families with errors introduced when the cDNA is first copied in the second strand reaction, where the error would be present in one of the first two DNA strands (Rate ~0.5). If an error occurs during the first cycle of PCR where both strands are copied, three of the four strands (rate ~0.75) would have the correct base. Errors in successive PCR cycles would result in greater agreement levels. Ignoring the frequency fluctuations due to stochastic template resampling, the 0.7 filter thus removes any families with 2^nd^ strand synthesis errors and/or which underwent sufficient recombination to drop agreement levels below 0.7. To help estimate the number of sUMI families which could fail this cutoff under different preparation conditions, the proportion of families containing < 30% reads which were identified as recombinant by UMI combinations are plotted in Fig. 2D. When no purification was performed, the fraction of sUMI families with < 30% recombinant reads decreased rapidly with increasing template input per PCR. In contrast, when using either of the two purification methods, the fraction of sUMI families with < 30% recombinant reads usually remained stable at ~ 95% until approximately 100 templates were added to the PCR. At template inputs > 100, fewer of the families from the bead-purified samples had < 30% recombinant reads compared to those purified by Blue Pippin, especially those with reduced cycle numbers, consistent with the results from the read quantitation shown in Fig. 2C. This analysis represents only the recombination observed by UMI combinations, thus in samples with low genetic diversity, recombination could take place with little impact on agreement or consensus sequence.

To help understand how the recombination observed in the UMI analysis could impact the recovery and accuracy of sequences calculated from sUMI compared to dUMI read collections, consensus sequences were generated using both methods. sUMI consensus are derived from all reads in a UMI-1 family, while only the reads from the most frequent UMI combination (rank 1) are used to create the corresponding dUMI consensus sequence. Aside from errors introduced during 2^nd^ strand synthesis, dUMI consensus sequences are therefore believed to be the most accurate. While Fig. 2D indicated the number of <u>families</u> estimated to pass the 0.7 filter based on prevalence of recombinant reads, the number of <u>sequences</u> from these families that were retained is shown in Fig. 2E. As predicted, many of the sUMI sequences from template inputs greater than 100 were discarded by the 0.7 filter, with unpurified samples suffering the greatest loss, however, not at levels as high as those predicted by the number of recombinant families. Presumably, as sample diversity increases, the proportion of sUMI sequences passing the 0.7 filter would fall to the levels predicted in Fig. 2D as the chance of recombining with a homologous molecule would go down with increasing diversity. As expected, nearly all dUMI sequences pass the 0.7 filter (mean 0.98 per sample, range 0.93-1). dUMI sequences that failed the 0.7 filter were investigated by aligning the reads, and in each site of discordance was a position with two bases present in similar frequencies, indicative of an early cycle PCR error, or a site of homopolymer length variation across the reads. These types of errors cannot be corrected for using either dUMI or sUMI methods.

Fig. 2F indicates the proportion of reads remaining after filtering. Because sUMI families contain reads with rank > 1, while dUMI families do not, the greater number of reads retained in the sUMI fraction represent rank >1 reads that do not cause the sUMI family to fail the 0.7 filter. This reinforces the hypothesis that with lower diversity many recombinant reads will have no effect on disagreement or the 0.7 filter, while greater diversity could reduce the proportion of accepted sUMI reads to levels similar to those found using dUMI.

When evaluating matched sUMI-dUMI consensus sequences from all preparation types, an error rate of 1 in ~339,000 bp was observed, hence 1 in every 156 of the 3kb sUMI sequences had a difference relative to the matched dUMI sequence. Samples with fewer than 100 templates that were also purified offered the lowest per-sequence and per-base error rates (Table 2). Seven families had discordant sUMI and dUMI sequences in samples with <100 input templates. The causes of discordance were investigated by aligning the reads from each family, then observing the nucleotide frequencies at the sites of disagreement and nucleotide linkage to each UMI-2 sequence. In all cases, discordance appeared to be caused by UMI collisions (i.e., two different cDNA molecules receiving the same UMI sequence by chance) or an error during 2^nd^ strand synthesis resulting in sites with similar nucleotide frequencies. From reactions with >100 templates, 25/34 families with discordant sUMI and dUMI sequences appeared to result from PCR recombination instead of UMI collisions or errors during 2^nd^ strand synthesis. In these families, reads with the most prevalent UMI-2 sequence were outnumbered by reads from all other UMI-2 sequences, which changed the majority nucleotide at that position. This demonstrates how artifactual consensus calls can occur when conditions allow high rates of PCR recombination.

**Table 2.**
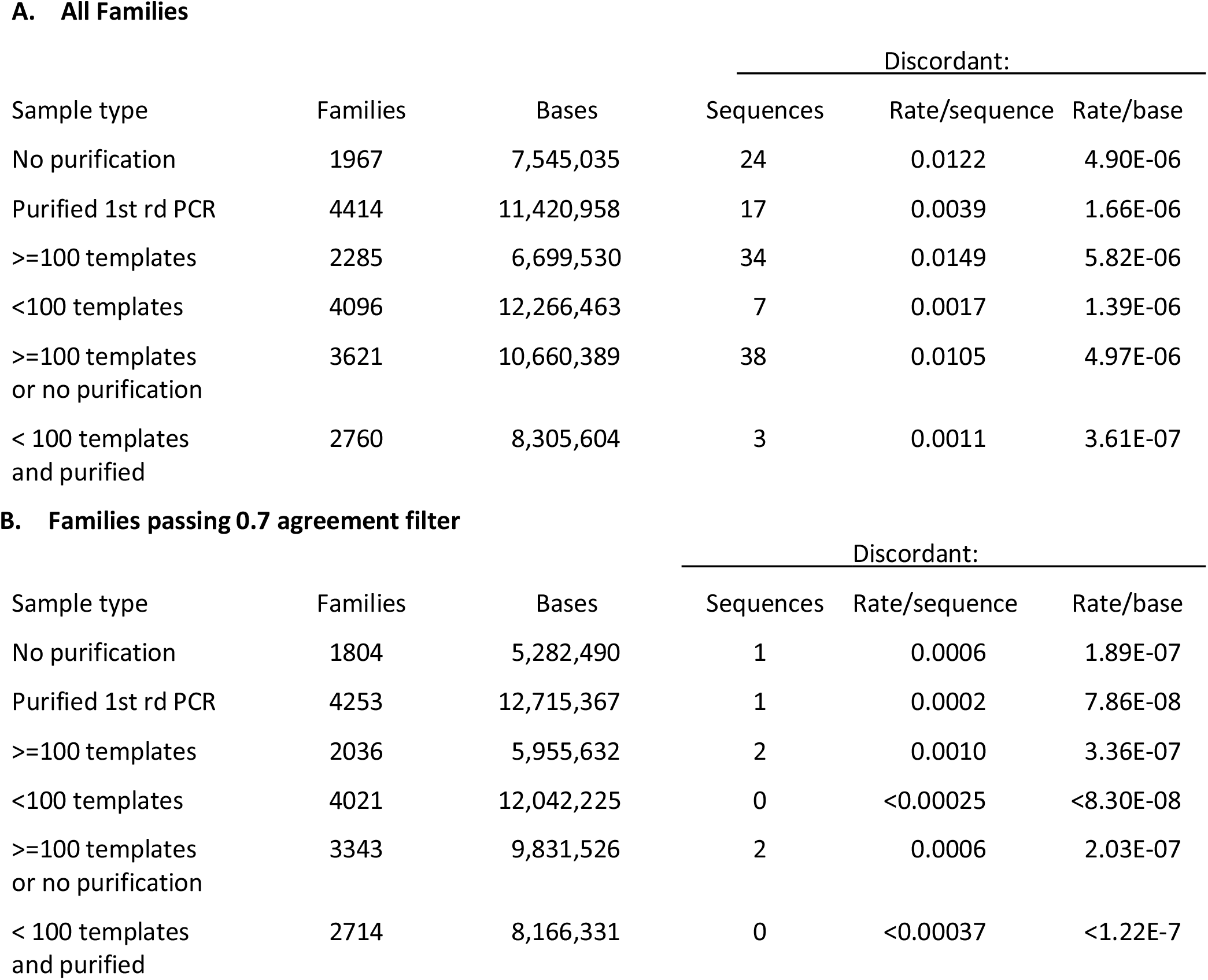
Discordant (error) rates using UMI and 0.7 minimum agreement filter.

Restricting the sUMI-dUMI sequence comparisons to all matched sequences passing the 0.7 filter, the per base error rate was reduced to 1 in ~1.1 × 10^7^ bp with only 1 in every 3,029 3kb sUMI sequences containing a difference. No differences were found between matched sUMI and dUMI sequences that passed the 0.7 filter in samples with fewer than 100 templates in the PCR (Table 2). This emphasizes the critical roll the 0.7 filter plays in reducing errors when utilizing the sUMI method.

## Discussion

In an effort to identify sUMI sample preparation methods allowing PCR amplification of diverse sequence populations while removing point mutational errors and without the formation of significant levels of artefactual recombinant molecules, we used a dUMI approach to track recombination events by differing UMI combinations and a novel pipeline to produce matched sUMI and dUMI consensus sequences. Relative to unpurified products, size selection using either bead or Blue Pippin purification after the first round of PCR resulted in greater recovery of the target amplicon, reduction of recombinant reads, and greater proportions of sequences passing 0.7 agreement filters, with the Blue Pippin method providing the best results. By reducing the number of cycles of the 2^nd^ round PCR by only 2, Blue Pippin purified samples with up to 277 templates per PCR resulted in fewer than 7% identifiably recombinant reads. This highlights the importance of using as few cycles as possible to generate sufficient DNA for sequencing, as well as the ability to add more templates if reducing cycle numbers. While reduced PCR cycles were not tested for all conditions, it can be reasonably assumed that reducing the number of cycles would have a similar effect on any preparation method. The only drawback to the use of fewer cycles is that more PCR reactions are likely to be required to obtain sufficient material for sequencing. While overall error rates were low when comparing sequences from sUMI and dUMI preparations, using fewer than 100 templates per PCR and either of the purification methods gave the lowest per base error rate. Ultimately, preparation methods cannot eliminate all recombination events and so sUMI families must also pass the 0.7 filter. When applying the 0.7 filter, all sequences from samples with less than 100 templates per PCR were identical to the corresponding dUMI sequences, allowing use of only a single UMI to maximize template recovery, and save time and reagent expense. The 0.7 filter, along with many additional features for analyzing sequences prepared with the sUMI approach, such as summary tables and plots, as well as phylogenetic trees (Fig. 3), have been packaged together as the PORPIDpipeline.

**Figure 3.**
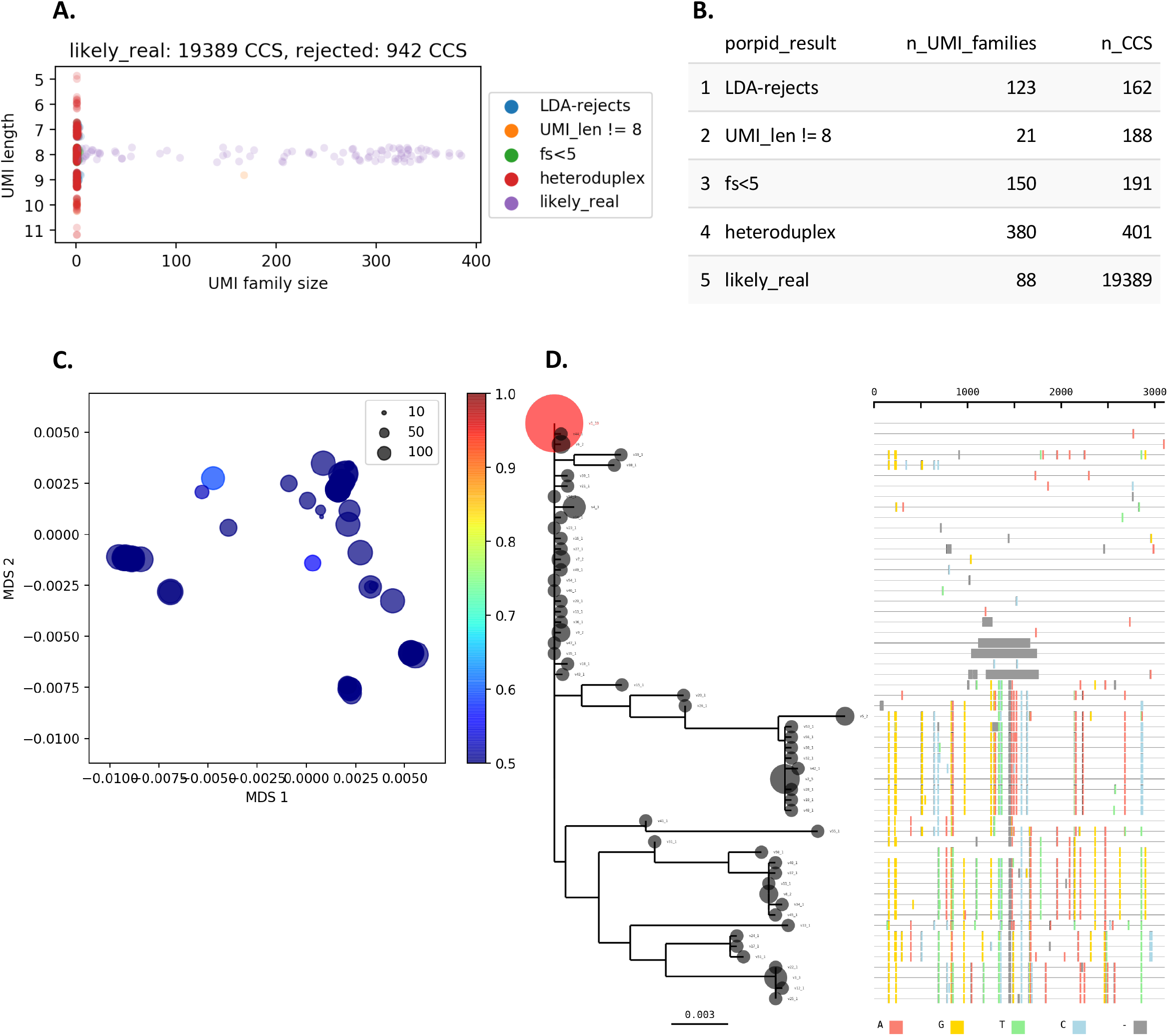
Elements of a PORPIDpipeline report. Panels A and B show elements of the results of PORPID processing, panels C and D show results of post processing steps of single-template consensus sequences. **A**. UMI stripplot. UMI family size (the number of individual CCS reads with a given UMI) is shown as a function of the UMI length determined by the PORPIDpipeline. Only “likely_real” UMIs are kept for downstream analysis. Those removed include the “LDA-rejects” that are likely generated by errors in the UMI sequence, “fs<5”, which indicates sequencing depth was under 5 CCS and considered too low for consensus analysis, and UMIs flagged as “heteroduplex” when the set of reads had a signature drop in quality in the UMI region indicative of a superimposed signal of two different UMI sequences during circular consensus generation. Some UMIs of length other than 8 pass all other criteria, but are still excluded from analysis (“UMI_len != 8”). **B**. UMI family and CCS totals are provided for each type of PORPID result post-processing of single-template consensus sequences. **C**. Multidimensional scaling (MDS) of template sequences and G>A APOBEC model. Classical MDS was used to represent all pairwise distances between template sequences in 2D space. Individual points are scaled by their family size (see inset key). A Bayesian model for APOBEC hypermutation is run on the single nucleotide substitution matrix between the global consensus and query sequence, and estimates the overall mutation rate and a G>A accelerator parameter from the data. Points are colored by probability that the G>A accelerator parameter is > 1. **D**. Phylogeny and highlighter plot of collapsed nucleotide variants. After collapsing by nucleotide sequence identity (Bubble size is scaled by the number of identical sequences), variants are numbered in descending order. The most frequent variant is used as the master sequence for the highlighter plot and the root for the phylogeny. Any variant above 10% of the population is colored in red on the tree. Colored lines in the highlighter plot represent different base substitutions or gaps (see key at the bottom of the panel).

In summary, limiting template input per PCR to 100, using the fewest PCR cycles which give sufficient material for sequencing, and performing a size selection after 1^st^ round PCR, allows for highly accurate sequence determination of large genomic fragments using a single UMI, PacBio SMRT sequencing, and PORPIDpipeline. This method allows for characterization of quasispecies at greater depths and gene lengths than previously possible by Sanger or short read sequencing methods and provides a streamlined and generalized sequencing approach which is highly adaptable to different pathogens and studies.

## Materials and Methods

### Specimen selection

Four plasma specimens derived from 4 individuals (A-D) from the RV217 clinical cohort[22, 23] were used for these studies (S1 Table). The RV217 study cohort enrolled seronegative individuals in East Africa and Thailand; participants were tested twice weekly with an HIV-1 RNA test. Plasma samples collected in the first days after HIV-1 diagnosis from three participants from Thailand and one participant from Kenya were selected. The protocol was approved by the Walter Reed Army Institute of Research and local ethics review boards: the Walter Reed Project, Kericho, Kenya and the Armed Forces Research Institute of Medical Sciences, Bangkok, Thailand. Only adult participants were enrolled. Written informed consent was obtained from all participants. Institutional Review Board approval was obtained for use of the samples, and all were anonymized.

### UMI Primer Composition

cDNA synthesis primers consisted of 1^st^ round reverse primer binding sequence (PB-R1-alt1 CCCGCGTGGCCTCCTGAATTAT), 2^nd^ round reverse primer binding sequence (PB-R2-alt1 CCGCTCCGTCCGACGACTCACTATA), 6-bp Sample ID sequence, 8-bp random UMI sequence, and a template binding sequence (Figure 1A). Second strand synthesis primers consisted of the 1^st^ round forward primer binding sequence (illu_F1 – AATGATACGGCGACCACCGA), 2^nd^ round forward primer binding sequence (illu_F2 – GATCTACACTCTTTCCCTACACG), 8-bp random UMI sequence, and a template binding sequence. All primers were synthesized by Integrated DNA Technologies and those containing a UMI sequence were synthesized with the “Hand-Mix” option selected for the 8-bp random UMI sequence. Subsequently, use of primers without the Hand-Mix option were validated. To allow pooling of multiple samples during sequencing, a different 6-bp Sample ID was used for each cDNA primer. Primer sequences and usage are listed in S2 Table.

### Library Amplification and Sequencing

RNA was extracted from plasma using the QIAamp Viral RNA Mini Kit (QIAGEN) and then used as template for cDNA synthesis, PCR and sequencing. 50 µl cDNA synthesis reactions were performed using Superscript III Reverse Transcriptase (Thermo Fisher Scientific) supplemented with 1U ThermaStop-RT (Sigma-Aldrich) per 50U SSIII to increases RT-PCR specificity. One µl of SSIII was added after 1.5 hours and incubated for another 1.5 hours to increase recovery of rare templates, however, similar results were obtained using SSIV with a 1 hour incubation. RNA was then removed by incubation with 2U RNase H [2U/µl, Invitrogen] for 20 min at 37°C. Each cDNA synthesis reaction utilized UMI-containing primers with a sample ID. To remove unincorporated primers, cDNA was purified with RNAClean XP magnetic beads (Beckman Coulter) at 1x ratio with 3x 80% ethanol washes and eluted into 25 µl of nuclease free water. Twenty-five µl of purified cDNA was subjected to 2^nd^ strand cDNA synthesis in a 40 µl reaction using 0.8 µl PrimeSTAR GXL (Takara Bio), 8 µl 5x Buffer, 3.2 µl dNTP Mix (2.5mM), and 10 pmol of a 2^nd^ strand primer mix containing UMI sequences. Reaction conditions were: 2 min at 98°C, 30 sec at 60°C, 10 min at 68°C and a 4°C hold. Second strand product was purified with AMpure XP magnetic beads in the same manner as the cDNA and eluted into 40 µl of nuclease free water.

The concentration of amplifiable templates in purified cDNA or 2^nd^ strand products was estimated by performing 25 µl nested PCRs with 0.5 µl PrimeSTAR GXL, 5 µl 5x Buffer, 2 µl dNTP Mix (2.5mM), and 10pmol of forward and reverse primers on limiting dilutions of the products then inputting the number of positive PCR per dilution to the web program Quality [https://indra.mullins.microbiol.washington.edu/quality/][24]. First round PCR cycling conditions (35 cycles) were: 2 min at 98°C; 35 cycles of 10 sec at 98°C, 15 sec at 60°C, and 1 min/kb extension at 68°C; followed by a 7 min extension at 68°C and a 4°C hold. Second round PCR cycling was identical except for using an annealing temperature of 62°C instead of 60°C. The pNL4-3 plasmid [25] was used as template in a positive control reaction with a template specific reverse primer spiked in, as the plasmid does not contain the primer binding sites incorporated by the UMI containing primers.

Using the estimated copy number results from Quality, 25 µl 1^st^ round PCRs were set up with an estimated 25, 50, or 100 UMI-labeled templates per reaction for Participants A, B, and C. Subject D had only enough 2^nd^ strand product for 25 templates per reaction. The number of reactions varied by sample from 1-10 with the goal sequencing at least 100 total templates per preparation. First round amplification conditions were the same as described for the template estimation reactions except that only 20 cycles of amplification were performed.

First round PCR products were pooled and mixed. 20 µl of each pool was diluted in 40 µl of 10 mM Tris-Cl, pH 8.0 to normalize across samples and ensure sufficient volume to test each treatment. Thirty µl of this volume was purified by size selection on a Blue Pippin instrument (Sage Science) using a 0.75% agarose cassette with Low Voltage 1-6kb definition and marker S1 on the Tight mode setting, using the size of the 1^st^ round amplicon as the Target. In parallel, 20 µl of the diluted volume was purified with AMpure XP magnetic beads at 0.5x ratio with 3x 80% EtOH washes and eluted into 26.7 µl of 10 mM Tris-Cl, pH 8.0. Lastly, 5 µl of the pooled PCR was diluted with 1.67 µl 10 mM Tris-Cl, pH 8.0 to match the dilution effects of the other methods.

One µl of the Blue Pippin, Ampure Bead or unpurified fractions was used as template in a 25 µl second round PCR. Conditions were the same as described for the template estimation reactions except that 20-22 cycles of amplification were performed. Blue Pippin purified fractions from the 100 estimated copy samples were also used as template in an additional 2^nd^ round PCR with only 20 cycles of amplification to test the effects of reducing cycle numbers. Twenty µl of each second round PCR reaction was purified with AMpure XP magnetic beads at 0.6x ratio with 3x 80% EtOH washes and eluted into 20 µl of 10 mM Tris-Cl, pH 8.0.

Since each sample from the same subject contained the same Sample ID, an additional round of PCR was performed to incorporate a unique pair of Index primers for each sample. 5 µl of purified second round PCR product was used as template in a 50 µl PCR with PrimeSTAR GXL containing a unique combination of one forward and one reverse Index primer (S2 Table). PCR conditions were the same as described above but with an altered cycling protocol: 2 min at 98°C; 5 cycles of 10 sec at 98°C, 30 sec at 55°C, and 1 min/kb extension at 68°C; followed by a 7 min extension at 68°C and a 4°C hold. PCR products were purified with AMpure XP magnetic beads at 0.6x ratio with 3x 80% EtOH washes and eluted into 40 µl of 10mM Tris-Cl, pH 8.0. If index primers were not necessary, a single cycle of synthesis and extension with fresh ingredients was performed to prevent heteroduplex formation. For this 5 µl of the following mixture was spiked into each 2^nd^ round PCR, 0.5 µl PrimeSTAR GXL, 1 µl 5x Buffer, 2 µl dNTP Mix (2.5mM), and 10pmol of forward and reverse primers, and incubate 2 min at 98°C, 15 sec at 62°C, and 10 minutes at 68°C.

Concentrations of purified DNA were determined using a Qubit dsDNA HS Assay Kit (Thermo Fisher). The samples were combined in equimolar amounts and the pool of samples was purified with AMpure XP magnetic beads at 0.7x ratio with 3x 80% EtOH washes and eluted into 40 µl of 10mM Tris-Cl, pH 8.0. The concentration of this pool along with other pools to be sequenced on the same SMRT cell was determined using Qubit. Library preparation was performed on each pool using the SMRTbell Express Template Prep Kit 2.0 (Pacific Biosciences) with each pool receiving a different barcoded adapter from the Barcoded Overhang Adapter Kit–8A (Pacific Biosciences). Completed libraries were purified with SMRTbell enzyme cleanup kit to remove incomplete or damaged SMRTbell molecules and then sequenced on a SMRT Cell 8M 15-hour movie using the Sequel II instrument (Pacific Biosciences).

### Bioinformatic Processing

Three snakemake[26] pipelines were created; one that performed the basic function of demultiplexing by Index primer (chunked_demux), one that performed additional demultiplexing by sample ID, UMI identification, and consensus sequence generation for sUMI and dUMI read collections (sUMI_dUMI_comparison) to test the different preparation methods, and one for use with the optimized sUMI approach with additional functionality after the consensus step (PORPIDpipeline). Because the sample preparations were labeled with index primers, the additional demultiplexing step (chunked_demux) was necessary to bin the reads into separate fastq collections by index combination and remove the index primer sequences. These collections were then used as input for the sUMI_dUMI_comparison pipeline where standard demultiplexing using the Sample ID took place.

All three pipelines were built with the Snakemake[26] workflow management system and provided as open source via Github [27, 28] [29]. Snakemake decomposes the workflow into rules that define how to obtain the stated output files from the given input files. The Snakefile lists the input and output files for each rule as well user defined parameters and the shell script that executes this process. Additionally, a user supplied config file lists each sample to be processed, along with each sample’s corresponding primer sequences and/or reference panels. A description of each rule in PORPIDpipeline is shown in Fig. 4 and described in more detail below. Only the early versions of the demux, porpid, and consensus rules were present in the sUMI_dUMI_comparison pipeline.

**Figure 4.**
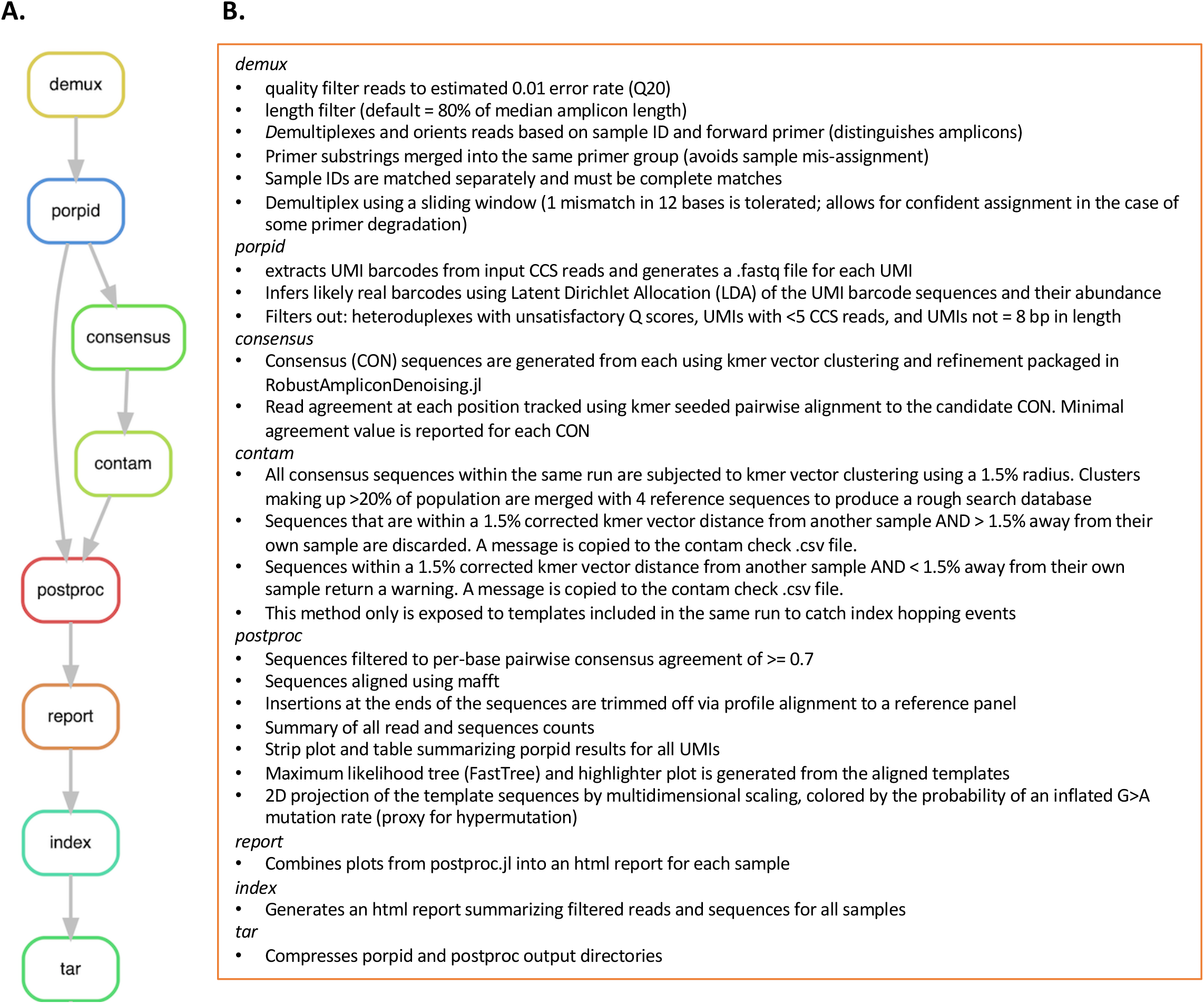
Overview of PORPIDpipeline. **A**. Flowchart illustrating flow between each rule within PORPIDpipeline and a brief description of each in **B**. See Methods for detailed descriptions. Only rules demux, porpid, and consensus were present in the pipeline used for analysis of the dUMI preparations.

#### Demux

*Quality Filtering and Demultiplexing*. After sequencing, PacBio raw reads are processed to produce circular consensus sequences (CCS), which are provided to the pipeline as fastq files (optionally gzipped). Each CCS is first examined and rejected if the mean error rate, inferred from the fastq quality scores[20], is greater than the user supplied parameter (default 0.01). Any CCS with length above or below user supplied parameters are also rejected. Each CCS is then assigned to a sample according to the PCR primers and Sample IDs listed in the config file, tolerating end degradation, and some imperfect matching on the primers, but requiring complete matching on the Sample IDs.

#### Porpid

*<u>P</u>robabilistic <u>O</u>ffspring <u>R</u>esolver for <u>P</u>rimer <u>ID</u>s*. For each sample, PORPID uses an alignment with the primer sequence to identify the UMI barcode in each CCS sequence, and groups sequences, by UMI, into “UMI families”. Since sequencing errors in the UMI itself can create erroneous “offspring” UMI barcodes, PORPID uses a probabilistic model, based on latent Dirichlet allocation (LDA) that considers the probability of generating each observed UMI from each other UMI by error, and then infers the underlying UMI frequencies from the noisily observed UMIs. UMI families that are likely to be erroneous, which are typically low-frequency neighbors of higher-frequency real UMI families, are excluded from downstream processing. Each UMI family is then examined and those that have UMI of length not equal to 8 or CCS number fewer than 5 are excluded. Next, UMIs produced as a result of heteroduplexes are removed. During PacBio SMRT sequencing, each strand of the circularized DNA template is successively sequenced. As the polymerase synthesizes each strand in a UMI library derived from a heteroduplex, two different UMI sequences will be read. This phenomenon produces a drop in PHRED scores[30, 31] for the UMI bases after circular consensus generation, since the two strands disagree, and we identify this disagreement using a combination of heuristic rules and statistical comparison to other regions. For the sUMI_dUMI_comparison pipeline, all UMI-2 sequences on the second strand were recorded, and not subjected to the filtering used for the UMI-1 sequence as described above (Table 1).

#### Consensus

*Consensus Sequence Generation*. Consensus sequences for UMI families that pass all quality control criteria are generated as described[32, 33]. Briefly, an exemplary read is selected based on the kmer distributions of each read, compared to the average for that UMI family, and then this is refined by alignment of all reads, modifying the consensus where the majority of reads mismatch. Read agreement to the final “polished” consensus is calculated and used downstream for the minimum agreement filter. The family size and minimum agreement values are reported for each consensus sequence and written as FASTA.

#### Contam

*Contamination Filter*. A rapid contamination screen is used, based on the observation that, for closely-related sequences, a kmer-derived distance well approximates an edit distance[32]. Consensus sequences for the same sample are subjected to kmer-vector clustering using a 1.5% radius. An overall centroid for each sample is merged with centroids for clusters making up more than 20% of the sample population and combined with kmer representations of common lab contaminants stored by the user in a supplied FASTA file. Default settings are such that sequences within a 1.5% distance from another cluster centroid and > 1.5% away from their own cluster centroid are discarded. Sequences that are within a 1.5% corrected kmer vector distance from another cluster centroid and < 1.5% away from their own cluster centroid are retained. Summary information is supplied to the user as a csv file.

#### Postproc

*Post-processing Steps*. For each sample, sequences that survived the contamination check are filtered using the minimum agreement values. This is a user supplied parameter but suggested to remain at default of 0.7. Insertions at the ends of the sequences are trimmed off via profile alignment to a user supplied reference panel. Major misalignments such as off target sequences (≥ 50% different from profile of a reference panel) are excluded. Outputs from postproc include the filtered sequence collection, mafft[34] alignment of these sequences, maximum likelihood tree, and a highlighter plot of collapsed sequences. Also produced are additional summary plots to be used by the Report rule.

#### Report

HTML reports are generated for each sample (Fig. 4). These contain a table summarizing rejected CCS, strip-plot indicating family sizes, maximum likelihood tree and highlighter plot of collapsed sequences, and a 2D projection of the sequences by multidimensional scaling, colored by the probability of an inflated G > A mutation rate using a modified nucleotide substitution model.

#### Index

An HTML index page is created containing summary tables and links to each sample report.

#### Tar

the porpid and postproc directories are archived and gzipped ready for downstream processing.

Compressed input/output files, config files, code written and executed in python and R studio, and sequence alignments were uploaded to the Dryad database upon submission. These files were used to collect and combine output files containing UMI information, calculate Levenshtein distance[35], perform recombination analyses, and write out data tables for plotting figures with Prism software (GraphPad Software). The python script was used to compare matched consensus sequence sets, and all alignments were created in Geneious software (BioMatters Ltd.). The README file and Analysis Flowchart (Fig. S2) indicate which files originate from each step and where they were used in the overall analysis.

## Author Contributions

D.H.W. and J.I.M. conceived of the laboratory methods, D.H.W. designed and performed optimization experiments. L.C. designed primers, and H.Z. performed experiments. W.D., A.P. and B.M. separately designed and evaluated steps in the bioinformatics pipeline with modifications converging onto the platform designed by B.M. H.M. refined the bioinformatics pipeline. M.R. provided the biological specimens for this study. C.W., B.M. and J.I.M. supervised different aspects of the work.

## Acknowledgements

We are indebted to the participants in the RV217 study and thank Merlin Robb, Leigh Anne Eller, Fred Sawe and Sorachai Nitayaphan. We thank Robert Ketteringham for contributions to the PORPID codebase and Lindsay Carpp for editorial assistance.

## Supporting information

**S1 Table.** Biological Samples. Thirty-three samples were generated from 4 four participants with varying viral loads by varying estimated input templates per PCR using QUALITY[24] (https://indra.mullins.microbiol.washington.edu/quality/), purification method of the 1st round PCR, and total cycle numbers. Also listed are the actual template numbers recovered after sequencing (sUMI families) and any PCR primers specific to a particular participant (primer sequences provided in S2 Table).

| Participant | RNA c/ml | cDNA Primer | Sample ID |
| --- | --- | --- | --- |
| A | 3,162,278 | PB_A5_R9013 | 5 |
| B | 1,737,801 | PB_A14_R9013 | 14 |
| C | 371,535 | PB_A9_R9013 | 9 |
| D | 1,023,293 | PB13_R9165 | 13 |

**B.**
| Sample | Estimated Templates in PCR | 1st Rd PCR Rxns | Purification Method | Total PCR Cycles | Actual Templates in PCR | Forward Index | Reverse Index |
| --- | --- | --- | --- | --- | --- | --- | --- |
| **Participant A** |  |  |  |  |  |  |  |
| 1 | 25 | 10 | None | 47 | 21 | F01 | R01 |
| 2 | 25 | 10 | Ampure XP Bead | 47 | 21 | F02 | R02 |
| 3 | 25 | 10 | Blue Pippin | 47 | 21 | F03 | R03 |
| 4 | 50 | 5 | None | 47 | 42 | F04 | R04 |
| 5 | 50 | 5 | Ampure XP Bead | 47 | 42 | F05 | R01 |
| 6 | 50 | 5 | Blue Pippin | 47 | 42 | F06 | R02 |
| 7 | 100 | 2 | None | 47 | 98 | F07 | R03 |
| 8 | 100 | 2 | Ampure XP Bead | 47 | 98 | F08 | R04 |
| 9 | 100 | 2 | Blue Pippin | 47 | 98 | F09 | R01 |
| 10 | 100 | 2 | Blue Pippin | 45 | 98 | F10 | R02 |
| **Participant B** |  |  |  |  |  |  |  |
| 11 | 25 | 4 | None | 47 | 65 | F01 | R01 |
| 12 | 25 | 4 | Ampure XP Bead | 47 | 65 | F02 | R02 |
| 13 | 25 | 4 | Blue Pippin | 47 | 65 | F03 | R03 |
| 14 | 50 | 2 | None | 47 | 126 | F04 | R04 |
| 15 | 50 | 2 | Ampure XP Bead | 47 | 126 | F05 | R01 |
| 16 | 50 | 2 | Blue Pippin | 47 | 126 | F06 | R02 |
| 17 | 100 | 1 | None | 47 | 278 | F07 | R03 |
| 18 | 100 | 1 | Ampure XP Bead | 47 | 278 | F08 | R04 |
| 19 | 100 | 1 | Blue Pippin | 47 | 278 | F09 | R01 |
| 20 | 100 | 1 | Blue Pippin | 45 | 278 | F10 | R02 |
| **Participant C** |  |  |  |  |  |  |  |
| 21 | 25 | 4 | None | 47 | 35 | F01 | R01 |
| 22 | 25 | 4 | Ampure XP Bead | 47 | 35 | F02 | R02 |
| 23 | 25 | 4 | Blue Pippin | 47 | 35 | F03 | R03 |
| 24 | 50 | 2 | None | 47 | 73 | F04 | R04 |
| 25 | 50 | 2 | Ampure XP Bead | 47 | 73 | F05 | R01 |
| 26 | 50 | 2 | Blue Pippin | 47 | 73 | F06 | R02 |
| 27 | 100 | 1 | None | 47 | 158 | F07 | R03 |
| 28 | 100 | 1 | Ampure XP Bead | 47 | 158 | F08 | R04 |
| 29 | 100 | 1 | Blue Pippin | 47 | 158 | F09 | R01 |
| 30 | 100 | 1 | Blue Pippin | 45 | 158 | F10 | R02 |
| **Participant D** |  |  |  |  |  |  |  |
| 28 | 25 | 5 | None | 47 | 32 | F01 | R01 |
| 29 | 25 | 5 | Ampure XP Bead | 47 | 32 | F02 | R02 |
| 30 | 25 | 5 | Blue Pippin | 47 | 32 | F03 | R03 |

**S2 Table.**
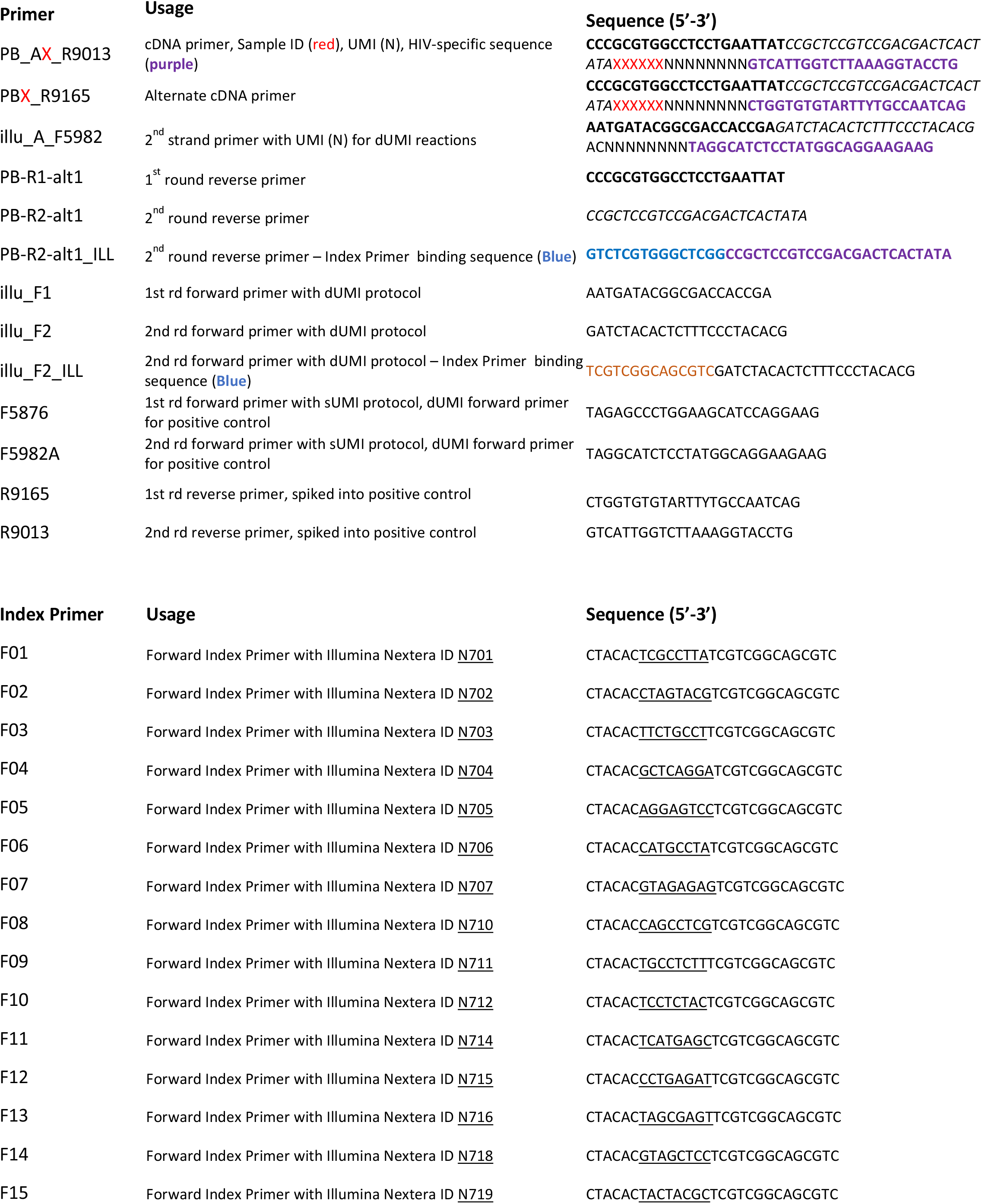
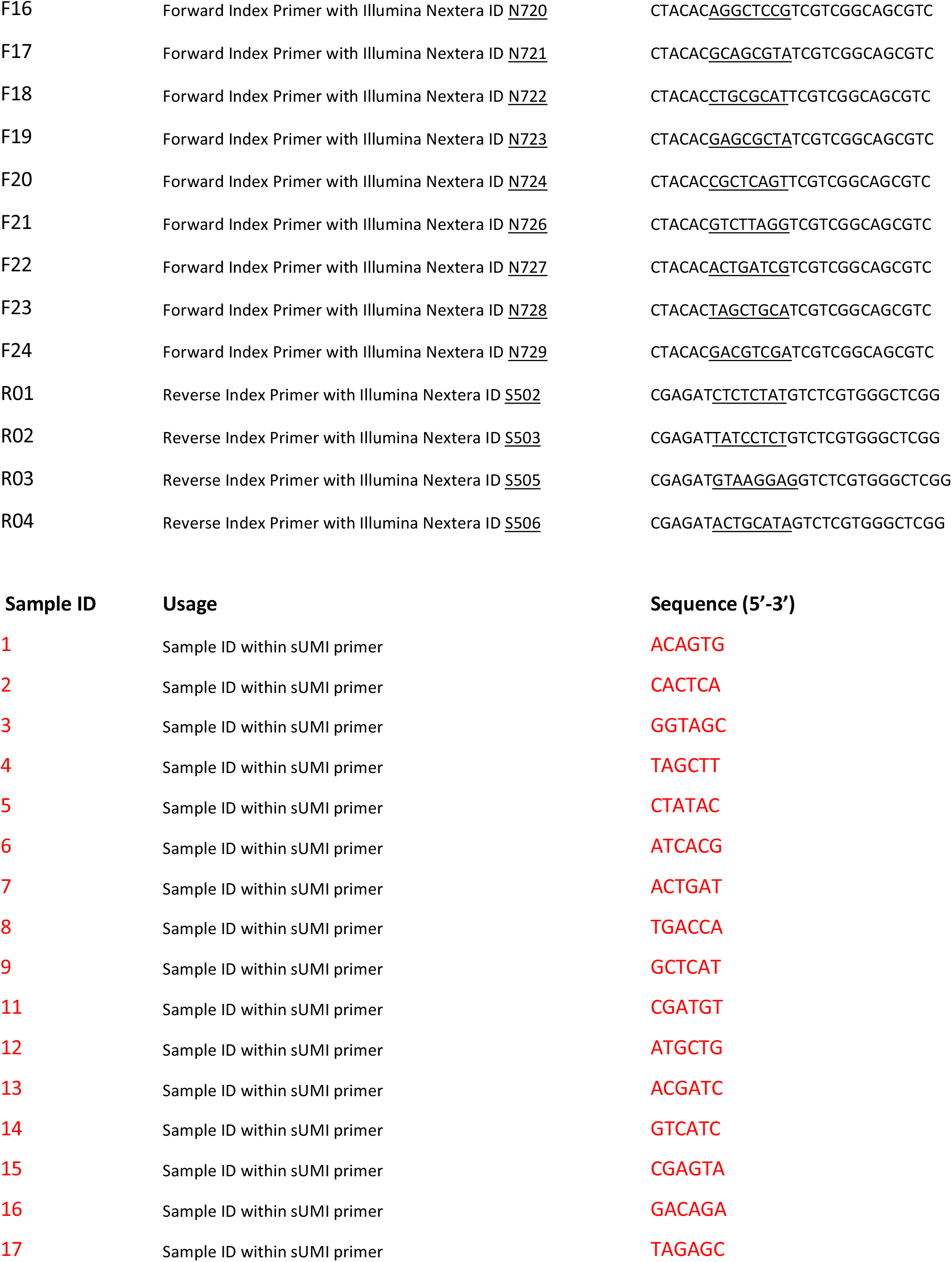
cDNA and PCR primer sequences. Name, usage, and sequence are listed for each primer. For each cDNA primer, one version is prepared with each of the 16 Sample ID sequences (indicated by XXXXXX).

**S1 Figure.**
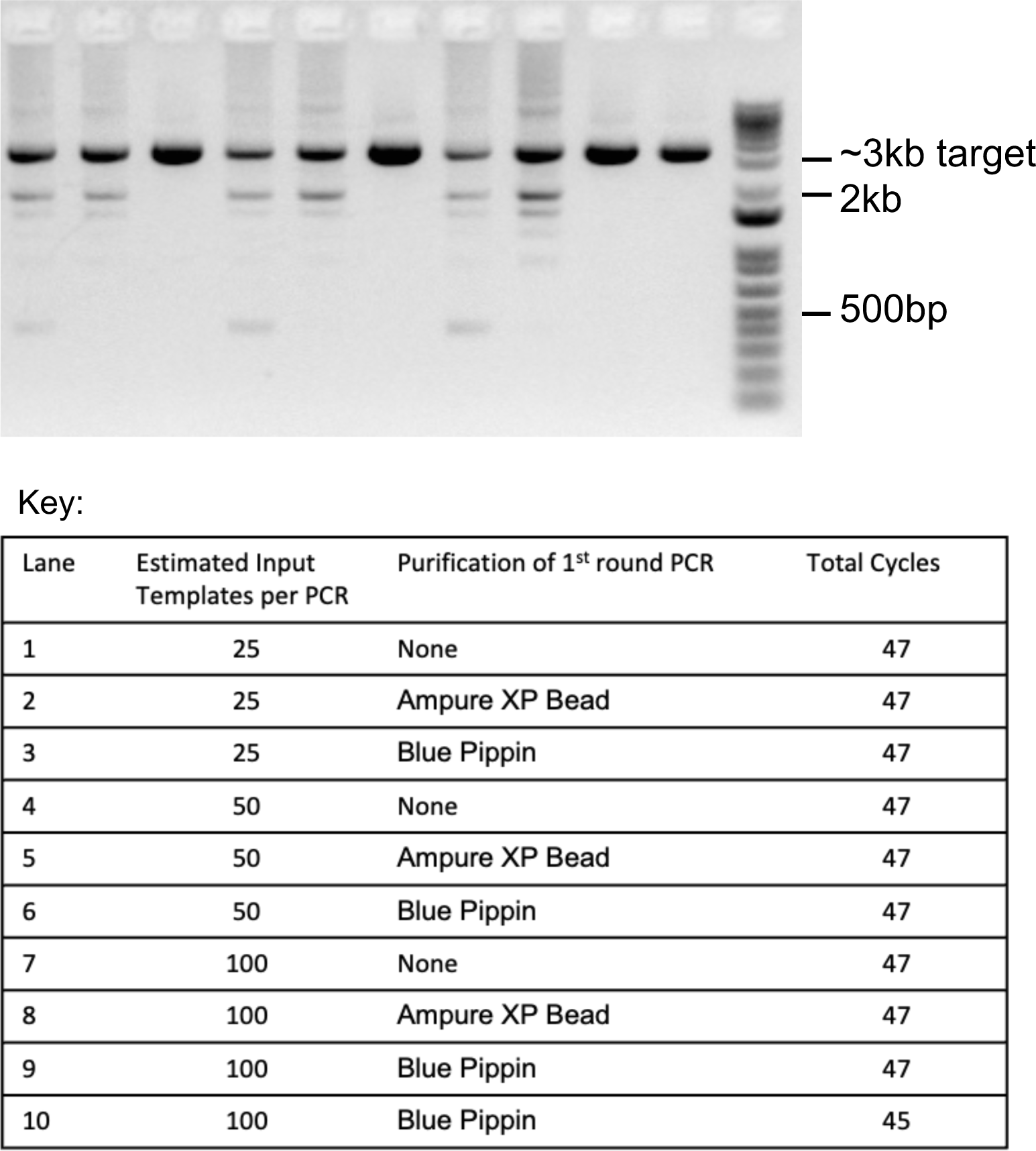
Gel electrophoresis and analysis of products according to purification method and PCR cycle number. 1% agarose gel image of purified PCR products after the 5 cycle Index PCR for different initial template inputs, methods of 1st round PCR purification, and total cycle numbers.

**S2 Figure.**
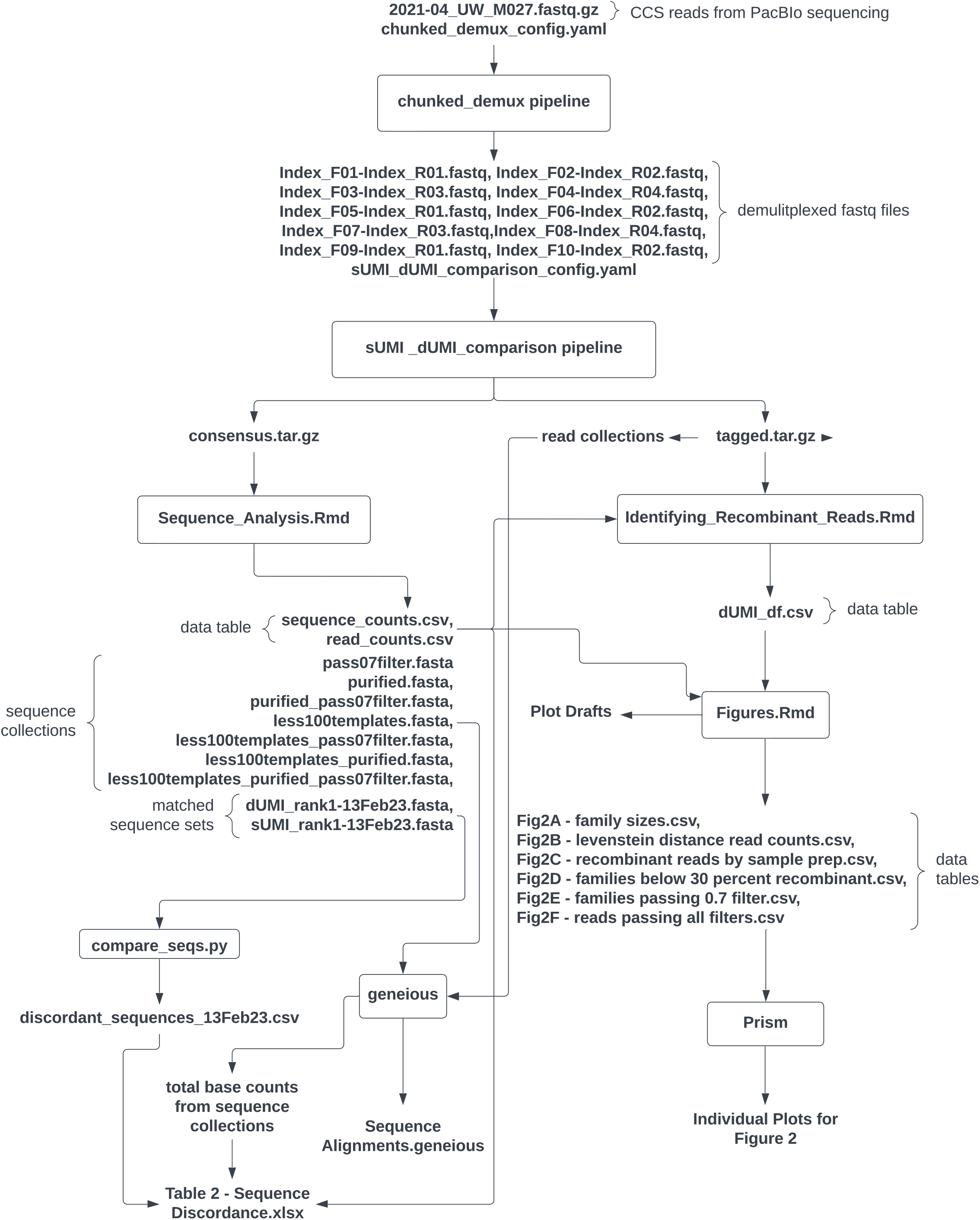
Analysis Flowchart. Beginning with the CCS read file from PacBio sequencing, the data moves through a series of pipelines, custom code, analysis steps, and software to create all tables and figures.

